# Fluctuations in population densities inform stability mechanisms in host-parasitoid interactions

**DOI:** 10.1101/2020.12.30.424820

**Authors:** Abhyudai Singh

## Abstract

Population dynamics of host-parasitoid interactions has been traditionally studied using a discrete-time formalism starting from the classical work of Nicholson and Bailey. It is well known that differences in parasitism risk among individual hosts can stabilize the otherwise unstable equilibrium of the Nicholson-Bailey model. Here, we consider a stochastic formulation of these discrete-time models, where the host reproduction is a random variable that varies from year to year and drives fluctuations in population densities. Interestingly, our analysis reveals that there exists an optimal level of heterogeneity in parasitism risk that minimizes the extent of fluctuations in the host population density. Intuitively, low variation in parasitism risk drives large fluctuations in the host population density as the system is on the edge of stability. In contrast, high variation in parasitism risk makes the host equilibrium sensitive to the host reproduction rate, also leading to large fluctuations in the population density. Further results show that the correlation between the adult host and parasitoid densities is high for the same year, and gradually decays to zero as one considers cross-species correlations across different years. We next consider an alternative mechanism of stabilizing host-parasitoid population dynamics based on a Type III functional response, where the parasitoid attack rate accelerates with increasing host density. Intriguingly, this nonlinear functional response makes qualitatively different correlation signatures than those seen with heterogeneity in parasitism risk. In particular, a Type III functional response leads to uncorrelated adult and parasitoid densities in the same year, but high cross-species correlation across successive years. In summary, these results argue that the cross-correlation function between population densities contains signatures for uncovering mechanisms that stabilize consumer-resource population dynamics.

## I. Introduction

Population dynamics of host-parasitoid interactions is typically formulated as a discrete-time model

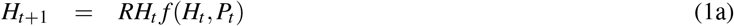

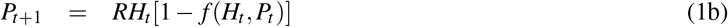

where *H_t_* and *P_t_* are the adult host, and the adult parasitoid densities, respectively, in year *t* [1]–[7]. Without loss of any generality, we assume that the host becomes vulnerable to parasitoid attacks in the larval stage. If *R* > 1 denotes the number of viable eggs produced by each adult host, then *RH_t_* is the host larval density exposed to parasitoid attacks. Adult (female) parasitoids search and attack host larvae with the density-dependent function *f*(*H_t_, P_t_*) < 1 representing the *escape response*, i.e., the fraction of host larvae escaping parasitism. Thus, *RH_t_f*(*H_t_, P_t_*) is the total larval density escaping parasitism that metamorphosize as adults the following year. Finally, *RH_t_* [1 – *f*(*H_t_, P_t_*)] is the density of parasitized larvae, where the juvenile parasitoid develops at the host’s expense by using it as a food source that ultimately results in host death. The juvenile parasitoids pupate and emerge as adult parasitoids the following year.

Perhaps the simplest formulation of (1) is the classical Nicholson-Bailey model

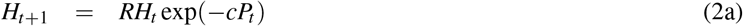

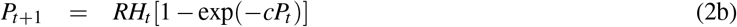

with a parasitoid-dependent escape response exp(-*cP_t_*), where *c* > 0 represents the rate at which parasitoids attack and parasitize host larvae [8]. The Nicholson-Bailey model is characterized by diverging oscillations in population densities resulting in an unstable population dynamics [8]. Much work has identified two orthogonal mechanisms by which stability can arise in these discrete-time models:

- The first mechanism is when the escape response *f*(*P_t_*) only depends on the parasitoid density, and then the non-trivial host-parasitoid equilibrium is stable, if and only, if, the equilibrium adult host density is an *increasing* function of the host reproduction rate *R* [9]. This type of stability arises through several related processes, such as, a fraction of the host population being in a refuge (i.e., protected from parasitoid attacks) [3], [10], large host-to-host difference in parasitism risk [9], [11]–[13], parasitoid interference [14]–[16], and aggregation in parasitoid attacks [17]–[19].
- The second mechanism is a Type III functional response where the parasitoid attack rate accelerates sufficiently rapidly with increasing host density [20]. Here the escape response *f* depends on both the host and parasitoid densities, and interestingly, in this case stability leads to the adult host equilibrium density being a *decreasing* function of the host reproduction rate *R* [21]. Parasitoids have tremendous potential for biological control of pest species [22]–[25], and a Type III functional response has been shown to suppress the host density to arbitrary low levels while maintaining system stability [21].

In this contribution, we consider annual variations in host reproduction that drive fluctuations in the host/parasitoid population densities. These random fluctuations are investigated in the context of two alternative stabilizing mechanisms: variation in parasitism risk across hosts and a Type III functional response. Our analysis develops analytical formulas that quantifies the extent of variations in population densities as a function of ecological parameters, and shows that harnessing the statistics of population fluctuations can be a vital tool for discriminating between stability mechanisms and characterizing host-parasitoid interactions. We start by reviewing how host-to-host differences in parasitism risk are incorporated in the Nicholson-Bailey model (2).

## II. Variation in parasitism Risk

The Nicholson-Bailey model assumes that all hosts are identical in terms of their vulnerability to parasitism. Perhaps a more realistic scenario is individual hosts differing in their risk of parasitism due to genetic factors, spatial heterogeneities, or are exposed to parasitoids for different durations, and at different times [26]–[29]. In essence, the attack rate *c* in (2) can be interpreted as “parasitism risk”, and by transforming it into a random variable we obtain

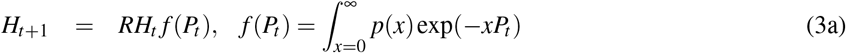

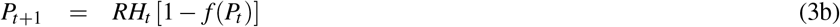

where *p*(*x*) is the distribution of risk across hosts [9]. A key assumption in this formulation is that risk is independent of the local host density, if hosts are non-uniformly distributed in space. Assuming *p*(*x*) follows a gamma distribution with mean 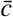 and coefficient of variation *CV* yields the escape response

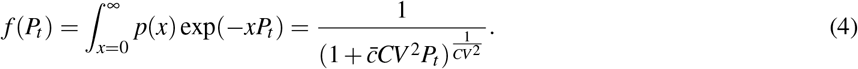

The non-trivial fixed point of the model (3)–(4) is given by

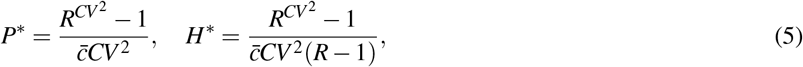

where *P*\* and *H*\* denote the parasitoid and host equilibrium densities, respectively. Prior analysis has shown that when the escape responsef *f*(*P_t_*) only depends on the parasitoid density, and then the non-trivial host-parasitoid equilibrium is stable, if and only, if,

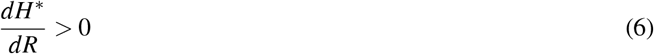

[9]. Applying this condition to (5) straightforwardly leads to a classical result - *CV* > 1 stabilizes the population dynamics irrespective of model parameters *R* and 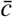 [11], [12], [19]. The stabilizing risk distribution implies that a majority of hosts are at low risk, and stability arises from parasitoid attacks being skewed towards a small fraction of high-risk individuals. This stability criterion motivated several studies investigating spatial patterns of parasitism in the field, and many data sets were found to be consistent with *CV* > 1 [13]. Recent work in this direction has relaxed the assumption of a gamma-distributed risk. It turns out that if *R* ≈ 1, then *CV* > 1 is the necessary and sufficient condition for stability irrespective of what form *p*(*x*) takes. However, for *R* ≫ 1, stability requires a skewed risk distribution with the modal risk being zero (as in the gamma distribution for *CV* > 1) [9].

## III. Incorporating yearly fluctuations in Host reproduction

Working with model (3)–(4) that considers a Gamma distributed risk, we incorporate random fluctuations in host reproduction by replacing *R* with an independent and identically distributed random variable *R_t_* with mean *R* and variance 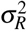. Considering small perturbations *h_t_*, *p_t_* around the equilibrium densities (5)

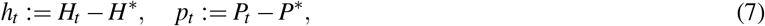

model (3)–(4) can be written as the following noise-driven linear discrete-time system

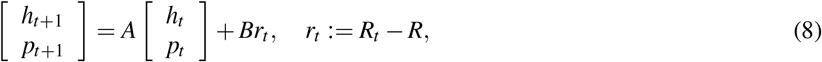

where the entries of the Jacobian matrix *A* are given by

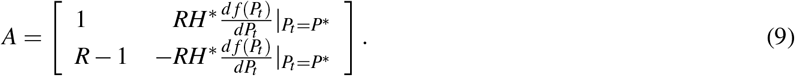

Here 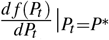 represents the derivative of the escape response with respect to *P_t_* evaluated at the equilibrium point, and assuming a stable host-parasitoid equilibrium, all eigenvalues of *A* are inside the unit circle [30]–[32]. The matrix *B* in (8) is given by

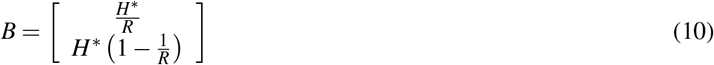

and characterizes the random forcing of the system by the zero-mean random variable *r_t_*.

Let

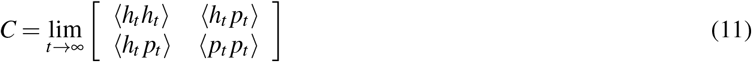

denote the steady-state covariance matrix, where 〈 〉 represented the expected value operation. Then the covariance matrix is the unique solution to the Lyapunov equation

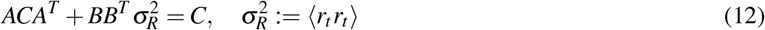

[33]. For a two-dimensional system, the Lyapunov equation can be solved analytically to yield

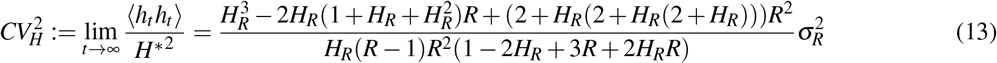

where 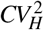 is the steady-state coefficient of variation squared of the host population density and

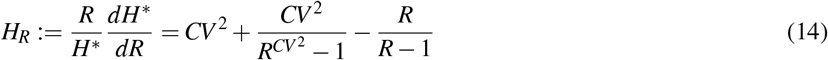

in the dimensionless log sensitivity of the host equilibrium density to *R*. Using (5), it can be seen that *H_R_* is monotonically related to the heterogeneity in the Gamma distributed risk as quantified by its coefficient of variation *CV*. Recall from (6) that the stability of the deterministic discrete-time system implies *H_R_* > 0. Interestingly, a close inspection of (13) reveals

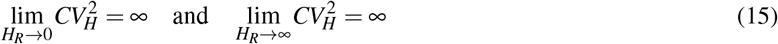

implying 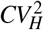 is minimized at an intermediate value of *H_R_*. For example, where *R* = 2, then (13) reduces to

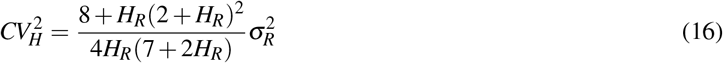

which is minimized where *H_R_* ≈ 1.51. From (14), this corresponds to host density fluctuations being minimal when *CV* ≈ 1.76 (Fig. 1). The magnitude of fluctuations in the parasitoid population density also follows a similar U-shape profile with increasing *CV*. Solving the Lyapunov equation (12) leads to the following Pearson correlation coefficient between the host and parasitoid densities (assuming *R* = 2)

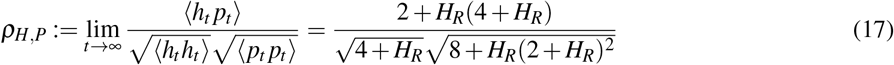

that predicts a moderate to strong correlation depending on *H_R_* (Fig. 2). This strong correlation can be intuitively explained by the fact that both the host and parasitoid equilibrium densities (5) are monotonically increasing functions of *R*.

**Fig. 1:**
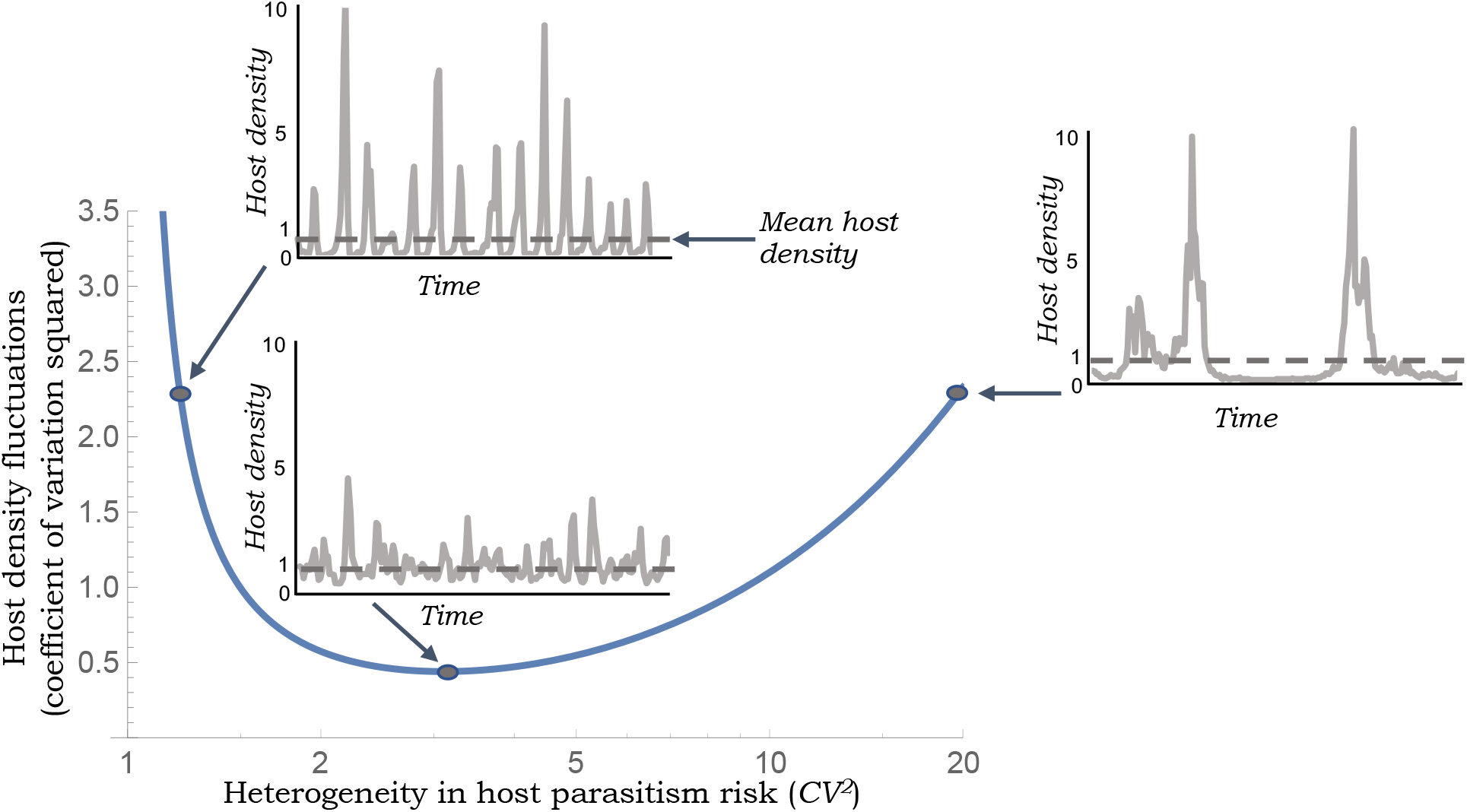
Extent of fluctuations in the host population density are minimized at an intermediate level of heterogeneity in parasitism risk. The steady-state coefficient of variation squared of the host population density as predicted by (14) and (16) is plotted as a function of the heterogeneity in parasitism risk *CV*. Three examples of host density fluctuations are generated by performing stochastic simulations of model (3)–(4) for different values of *CV* assuming 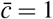, *R* = 2 and 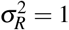. All time series are normalized to have a mean value of one.

**Fig. 2:**
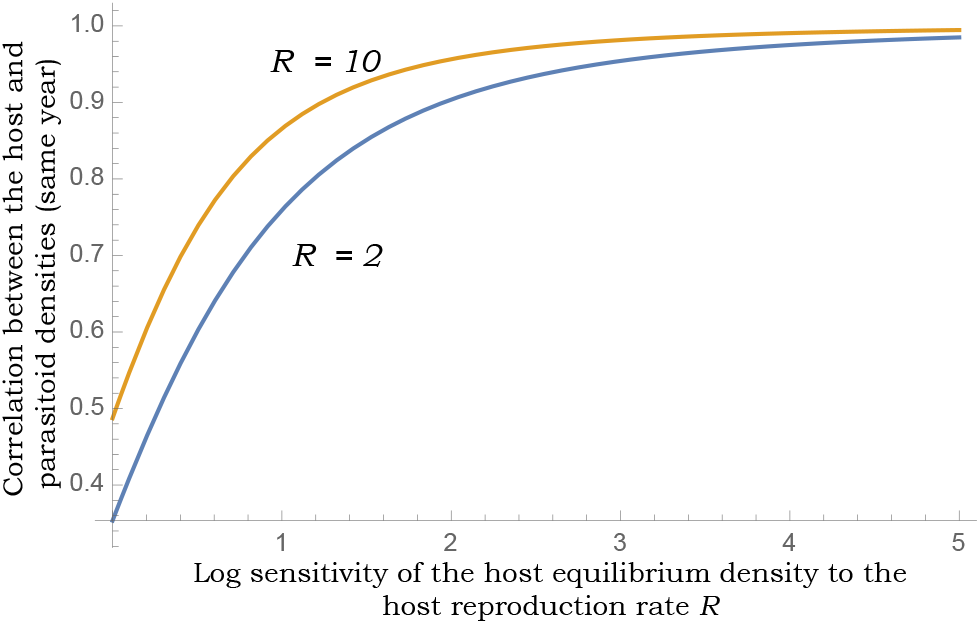
Randomness in host reproduction induces strong positive correlations between host-parasitoid densities for model (3)–(4) that incorporates heterogeneity in parasitism risk. Predicted Pearson correlation coefficient between the host and parasitoid densities for *R* = 2 and *R* = 10 as a function of *H_R_* as given in (14).

## IV. Stability arising through a Type III functional response

We next focus our attention to another stabilizing mechanism based on density-dependence in the parasitoid attack rate. In our prior work, we have considered a Type III parasitoid functional response, where the attack rate *cL^m^* accelerates with increasing host larvae density *L* for some positive constant *c* and exponent *m*. Here, *L* denotes the non-parasitized larval density that decreases over time during the vulnerable stage leading to a variable attack rate. To capture such effects of populations changing continuously within the larval stage of each year, a semi-discrete or hybrid formalism has been proposed to mechanistically formulate the corresponding discrete-time model. This semi-discrete approach relies on solving a continuous-time differential equation describing population interaction during the host’s vulnerable stage to derive update functions connecting population densities across consecutive years [20], [34]–[37]. For an attack rate *cL^m^* this leads to the model (1) with escape response

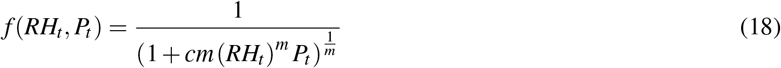

that depends on both host and parasitoid population densities [20]. It turns out that the model’s unique non-trivial fixed point

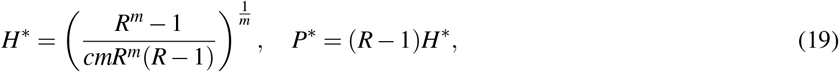

is stable iff *m* > 1, and *m* = 1 results in a neutrally stable equilibrium where populations oscillate with a period of 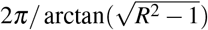 [20]. Interestingly, in contrast to (6), here *H*\* is a decreasing function of *R*, while *P*\* is an increasing function of *R*. It is important to point out that a phenomenological approach of incorporating a Type III functional response by simply substituting *c* in the Nicholson-bailey model (2) with *c*(*RH_t_*)*^m^* (i.e., the parasitoid attack rate is set by the initial larval density *RH_t_* and remains fixed through the larval stage) leads to an unstable population equilibrium for all *m* ≥ 0 [38], [39].

As done in the previous section, considering stochastic fluctuations in the host reproduction rate in the model defined by (1) and (18) yields the Lyapunov equation (12) with

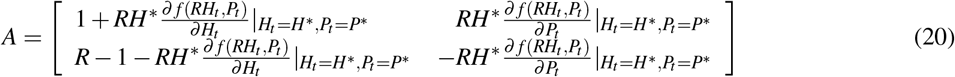

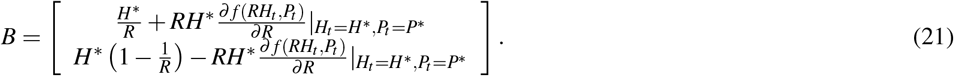

Solving the Lyapunov equation reveals that in this case the extent of fluctuations in host/parasitoid densities monotonically decreases to zero with increasing *m*. For example, for *R* = 2, 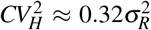 when *m* = 2 and 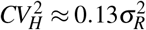 when *m* = 3. This makes sense as increasing *m* not only increases system stability (i.e., faster return to equilibrium in response to perturbations) but also makes the host equilibrium less sensitive to *R* with

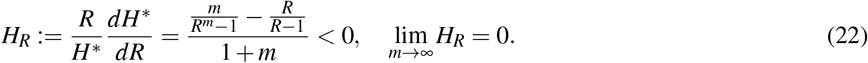

We also obtain the following analytical expression for the cross-species Pearson correlation (assuming *R* = 2)

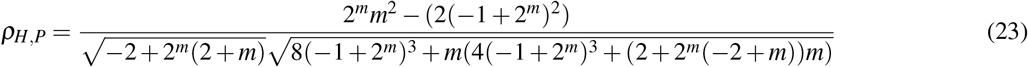

that is predicted to be *ρ_H,P_* ≈ −0.01 when *m* = 2 and *ρ_H,P_* ≈ −0.03 when *m* = 3. Such uncorrelated fluctuations in host/parasitoid densities in response to random perturbations in *R* is reflective of *H*\* and *P*\* in (19) being a decreasing and increasing function of *R*, respectively. Intriguingly, if one quantifies the cross-correlation function across different years, i.e., the correlation between *H_t_* and *P*_*t*+Δ*t*_ where Δ*t* is the generation lag of the host with respect to the parasitoid, then one sees a sharp jump to positive cross-species correlation between *H_t_* and *P*_*t*+1_ which then again goes back to zero with larger generation lags (Fig. 3). Hence, a Type III functional response is characterized by uncorrelated same-year fluctuations in population densities that exhibit a non-monotonic crosscorrelation function profile. In contrast to these results, heterogeneity in parasitism risk leads to strong same-year correlations that gradually decay to zero with increasing Δ*t* (Figs. 2 and 3). The qualitative differences in the cross-correlation function profile provides a valuable tool to discriminate between these two stabilizing mechanisms from time-series data.

**Fig. 3:**
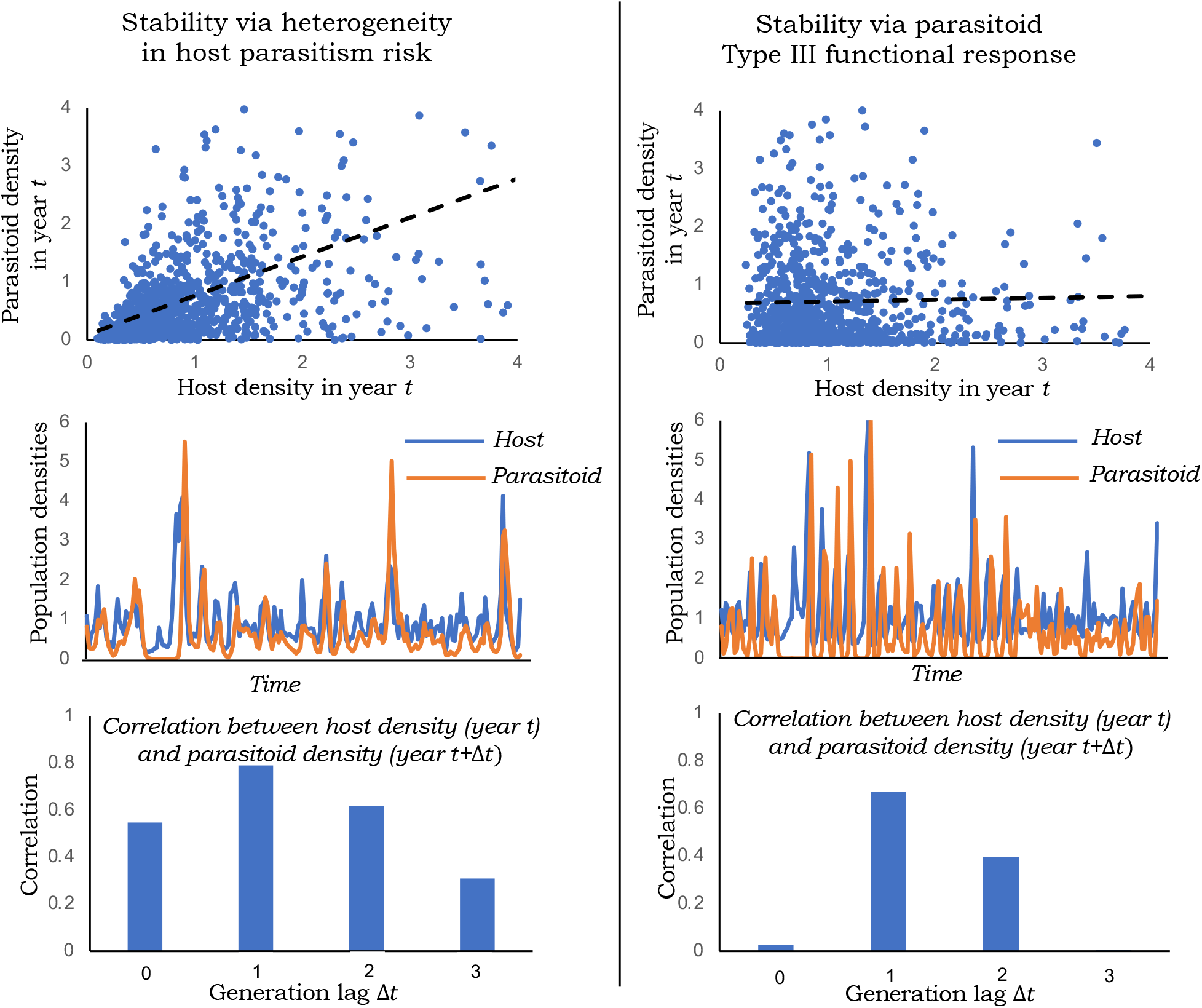
Different stabilizing mechanisms of host-parasitoid population dynamics can be discriminated from the cross-correlation function profile. *Left*: Scatter plot of *H_t_* and *P_t_* from a stochastic simulation of model (3)–(4) for *CV* = 2, 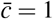, *R* = 2, 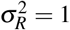 along with the time-series of host/parasitoid population densities. Heterogeneity in parasitism risk results in strong correlation between *H_t_* and *P_t_*, that shows a modest increase followed by decay to zero with increasing generation lag Δ*t* for correlations between *H_t_* and *P*_*t*+Δ*t*_. *Right*: Scatter plot of *H_t_* and *P_t_* from a stochastic simulation of model (1) and (18) for *m* = 2, 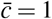, *R* = 2, 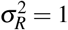 reveals uncorrelated fluctuations in host/parasitoid densities for a Type III functional response. As can be seen in the simulated time-series and the cross-correlation function, *h_t_* and *p*_*t*+1_ show a strong positive correlation that decays back to zero with increasing generation lag Δ*t*.

## References

[1] M. P. Hassell. New York: Oxford University Press, 2000.

[2] W. S. C. Gurney and R. M. Nisbet, Ecological Dynamics. Oxford University Press, 1998.

[3] W. W. Murdoch, C. J. Briggs, and R. M. Nisbet, Consumer-Resouse Dynamics. Princeton, NJ: Princeton University Press, 2003.

[4] N. Kakehashi, Y. Suzuki, and Y. Iwasa, “Niche overlap of parasitoids in host-parasitoid systems: its consequence to single versus multiple introduction controversy in biological control,” Journal of Applied Ecology, pp. 115–131, 1984.

[5] R. M. May and M. P. Hassell, “The dynamics of multiparasitoid-host interactions,” The American Naturalist, vol. 117, no. 3, pp. 234–261, 1981.

[6] E. Hackett-Jones, C. Cobbold, and A. White, “Coexistence of multiple parasitoids on a single host due to differences in parasitoid phenology,” Theoretical Ecology, vol. 2, no. 1, pp. 19–31, 2009.

[7] E. van Velzen, S. Pérez-Vila, and R. S. Etienne, “The role of within-host competition for coexistence in multiparasitoid-host systems,” The American Naturalist, vol. 187, no. 1, pp. 48–59, 2016.

[8] A. Nicholson and V. A. Bailey, “The balance of animal populations. part 1.” Proc. of Zoological Society of London, vol. 3, pp. 551–598, 1935.

[9] A. Singh, W. W. Murdoch, and R. M. Nisbet, “Skewed attacks, stability, and host suppression,” Ecology, vol. 90, no. 6, pp. 1679–1686, 2009.

[10] E. Bešo, S. Kalabušić, N. Mujić, and E. Pilav, “Stability of a certain class of a host–parasitoid models with a spatial refuge effect,” Journal of Biological Dynamics, vol. 14, no. 1, pp. 1–31, 2020.

[11] A. D. Taylor, “Heterogeneity in host-parasitoid interactions: ’aggregation of risk’ and the ’cv^2^>1 rule.’,” Trends in Ecology and Evolution, vol. 8, pp. 400–405, 1993.

[12] M. P. Hassell, R. M. May, S. W. Pacala, and P. L. Chesson., “The persistence of host–parasitoid associations in patchy environments. I. a general criterion.” American Naturalist, vol. 138, pp. 568–583, 1991.

[13] S. W. Pacala and M. P. Hassell., “The persistence of host– parasitoid associations in patchy environments. II. evaluation of field data.” American Naturalist, vol. 138, pp. 584–605, 1991.

[14] C. Bernstein, “Density dependence and the stability of host-parasitoid systems,” Oikos, pp. 176–180, 1986.

[15] C. Free, J. Beddington, and J. Lawton, “On the inadequacy of simple models of mutual interference for parasitism and predation,” The Journal of Animal Ecology, pp. 543–554, 1977.

[16] D. Rogers and M. Hassell, “General models for insect parasite and predator searching behaviour: interference,” The Journal of Animal Ecology, pp. 239–253, 1974.

[17] J. D. Reeve, J. T. Cronin, and D. R. Strong., “Parasitoid aggregation and the stabilization of a salt marsh host– parasitoid system,” Ecology, vol. 75, pp. 288–295, 1994.

[18] P. Rohani, H. C. J. Godfray, and M. P. Hassell, “Aggregation and the dynamics of host-parasitoid systems: A discrete-generation model with within-generation redistribution,” The American Naturalist, vol. 144, no. 3, pp. 491–509, 1994.

[19] R. M. May, “Host–parasitoid systems in patchy environments: a phenomenological model,” Journal of Animal Ecology, vol. 47, pp. 833–844, 1978.

[20] A. Singh and R. M. Nisbet, “Semi-discrete host-parasitoid models,” Journal of Theoretical Biology, vol. 247, no. 4, pp. 733–742, 2007.

[21] A. Singh and B. Emerick, “Hybrid systems modeling of ecological population dynamics,” bioRxiv, 2020.

[22] S. D. Lane, C. M. St. Mary, and W. M. Getz, “Coexistence of attack-limited parasitoids sequentially exploiting the same resource and its implications for biological control,” in Annales Zoologici Fennici. JSTOR, 2006, pp. 17–34.

[23] B. S. Pedersen and N. J. Mills, “Single vs. multiple introduction in biological control: the roles of parasitoid efficiency, antagonism and niche overlap,” Journal of Applied Ecology, vol. 41, no. 5, pp. 973–984, 2004.

[24] P. K. Abram, J. Brodeur, V. Burte, and G. Boivin, “Parasitoid-induced host egg abortion; an underappreciated component of biological control services provided by egg parasitoids.” Biological Control, no. 98, pp. 52–60, 2016.

[25] M. A. Jervis, B. A. Hawkin, and N. A. C. Kidd, “The usefulness of destructive host-feeding parasitoids in classical biological control: theory and observation conflict,” Ecological Entomology, vol. 21, no. 1, pp. 41–46, 1996.

[26] T. Okuyama, “Density-dependent distribution of parasitism risk among underground hosts.” Bulletin of Entomological Research, vol. 109, no. 4, pp. 528–533, 2019.

[27] C. A. Cobbold, J. Roland, and M. A. Lewis, “The impact of parasitoid emergence time on host-parastioid population dynamics,” Theor Popul Biol, vol. 75, no. 2, pp. 201–215, 2009.

[28] H. Liere, D. Jackson, and J. Vandermeer, “Ecological complexity in a coffee agroecosystem: spatial heterogeneity, popoulation persistence and biological control,” PLoS One, vol. 7, no. 9, 2012.

[29] N. Zoroa, E. Lesigne, M. J. Fernandez-Saez, P. Zoroa, and J. Casas, “The coupon collector urn model with unequal probabilities in ecology and evolution,” Journal of The Royal Society Interface, vol. 14, no. 127, 2017.

[30] G. Ledder, Mathematics for the life sciences: calculus, modeling, probability, and dynamical systems. Springer Science & Business Media, 2013.

[31] S. Elaydi, An Introduction to Difference Equations. Newyork: Springer, 1996.

[32] R. M. Nisbet and W. Gurney, Modelling fluctuating populations: reprint of first edition (1982), 2003.

[33] Z. Gajic and M. T. J. Qureshi, Lyapunov matrix equation in system stability and control. Courier Corporation, 2008.

[34] A. Singh and R. M. Nisbet, “Variation in risk in single-species discrete-time models,” Mathematical Biosciences and Engineering, vol. 5, pp. 859–875, 2008.

[35] B. K. Emerick and A. Singh, “The effects of host-feeding on stability of discrete-time host-parasitoid population dynamic models.” Mathematical Biosciences, vol. 272, pp. 54–63, 2016.

[36] E. Pachepsky, R. M. Nisbet, and W. W. Murdoch, “Between discrete and continuous: Consumer-resource dynamics with synchronized reproduction,” Ecology, vol. 89, no. 1, pp. 280–288, 2007.

[37] B. K. Emerick and A. Singh, “Global redistribution and local migration in semi-discrete host-parasitoid population dynamic models.” Mathematical Biosciences, vol. 327, p. 108409, 2020.

[38] D. J. Rogers, “Random searching and incest population models,” J. of Animal Ecology, vol. 41, pp. 369–383, 1972.

[39] M. P. Hassell and H. N. Comins, “Sigmoid functional responses and population stability,” Theoretical Population Biology, vol. 14, pp. 62–66, 1978.

